# Computational biophysical characterization of a superradiant virus-like particle in its ground state

**DOI:** 10.1101/2025.10.15.682673

**Authors:** Anjali Krishna, Anoushka Buddhikot, Jodi A. Hadden-Perilla

## Abstract

Superradiance (SR) arises when fluorophores radiate collectively rather than independently, producing ultrabright emission bursts. Recent experiments show that dye-labeled virus-like particles (VLPs) can exhibit SR at room temperature, where the VLP scaffold organizes dyes into a macrodipole of coherent emitters under pulsed excitation. To elucidate the biophysical basis of this phenomenon, microsecond-scale all-atom molecular dynamics (MD) simulations were performed on intact Brome mosaic virus (BMV) VLPs, comparing the native scaffold to a capsid decorated with Oregon Green 488 (OG). Results show that OG conjugation at K105 induces only subtle structural changes: a slight expansion of VLP volume and marginally higher inter-subunit flexibility, while overall shell morphology and pore architecture remain intact. However, dianionic OG alters VLP electrostatics, increasing water exchange across the shell and reducing chloride permeability. Dyes bound in pentamers are more closely packed and conformationally restricted than those in hexamers, and their transition dipoles more frequently adopt parallel orientations predicted to favor excitonic coupling and cooperative emission. By examining the role of scaffold architecture in dye interactions and dynamics, this work offers a mechanistic framework and design principles for engineering SR-VLPs as next-generation fluorescent probes and photonic materials.

## Introduction

Virus-like particles (VLPs) are nanostructures that self-assemble from viral proteins. They mimic the architecture of native viruses but lack genetic material and are non-infectious. Their symmetry, stability, and ease of chemical modification make them valuable tools in targeted gene and drug delivery, vaccine and materials design, and diagnostics.^1–3^ The plant pathogen Brome mosaic virus (BMV) is a major platform for functionalized VLP development.^4^ Recent studies have shown that covalent decoration of BMV with fluorescent dyes such as Oregon Green 488 (OG) can yield superradiant particles. ^5–7^ As the number of fluorophores per VLP approaches a critical threshold, an ultra-bright radiation occurs under pulsed excitation. The combination of increased emission intensity and accelerated decay is a hallmark of cooperative relaxation, suggesting that the dyes emit radiation collectively rather than independently.^5,7^ The spectral and temporal signatures of this emission are consistent with that of superradiance (SR). ^8^ Replacement of the VLP with spherical nanoparticles such as glass beads fails to yield SR,^5^ indicating that the symmetry and biophysical properties of BMV are critical to emergent optical phenomena but absent in other scaffolds.

Unlike conventional multi-fluorophore emitters that are limited by random, uncorrelated emission, self-quenching, and photobleaching, ^9^ dye-labeled BMV produces intense, synchronized SR emission with a high signal-to-noise ratio. Remarkably, despite limited previous examples of SR beyond liquid-helium-cooled conditions, ^10^ dye-labeled BMV exhibits SR even at room temperature. Thus, these SR particles hold promise for improving fluorescent imaging and photonics applications, including with respect to intraoperative cancer imaging,^11^ bioaerosol detection,^12^ and anti-counterfeiting.^13^ While SR appears to depend strongly on the spatial order and dynamics of the dye–protein system, where any disruption to VLP structure results in the abrogation of collective emission, ^5,7^ the detailed mechanism by which BMV enables coherent coupling of fluorophores remains unclear. The locations of reactive lysines^14^ to which dyes are covalently attached to the VLP appear to play a key role. ^6^

BMV forms stable VLPs that are easily assembled, relatively rigid, and tolerant of chemical modification,^14,15^ resulting in consistently high labeling yields with dyes like OG (Figure 1a, inset). The structural coat of BMV, called a capsid, is composed of 180 protein monomers, approximately 20 kDa each.^16^ These proteins adopt three quasi-equivalent conformations A, B, and C (Figure 1a), which together form the asymmetric unit of the 28-nm capsid (Figure 1b). The capsid exhibits T=3 icosahedral symmetry, organized as 12 pentamers and 20 hexamers (Figure 1c-d). Pentamers comprise five copies of the A chain, and hexamers comprise two copies each of the B and C chains. While residues 42–189 form the ordered capsid shell, residues 1-41 of the coat protein represent an intrinsically disordered N-terminal domain that extends into the interior of the virion and contributes to RNA encapsidation and inter-subunit contacts.^17^ Dye conjugation on the flexible N-termini does not contribute to SR,^6^ implicating labeling positions on the ordered portion of the shell.

**Figure 1:**
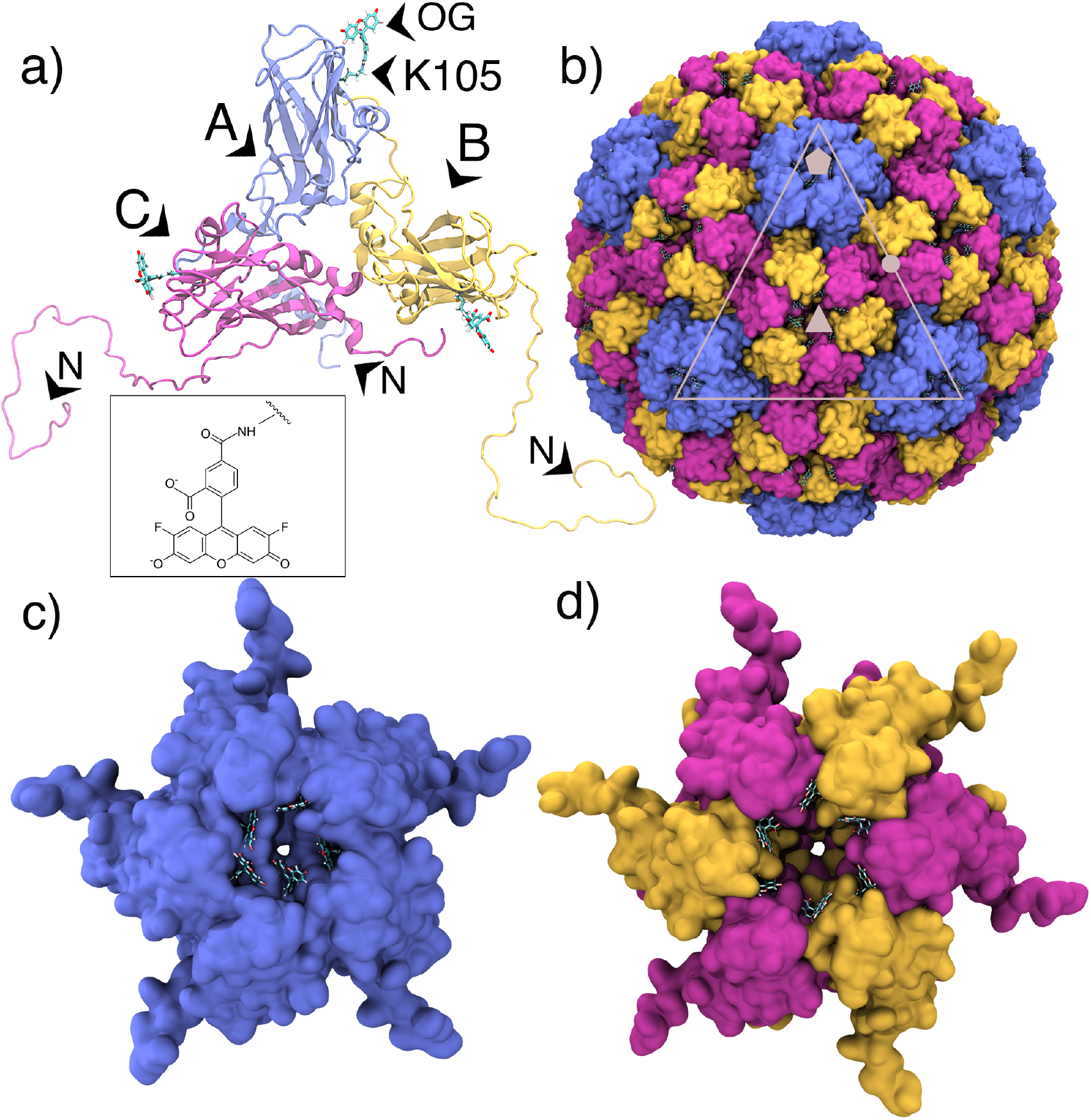
**(a)** Asymmetric unit of BMV capsid, composed of three quasi-equivalent chains A (purple), B (yellow), and C (magenta), each labeled with OG (cyan) at K105. OG is attached via an amide bond to the *E*-ammonium group of the lysine side chain. Inset: chemical structure of OG. **(b)** Surface representation of the intact T=3 capsid composed of 180 coat protein subunits, colored by subunit type (A, B, C) with icosahedral fivefold, threefold/quasi-sixfold and two-fold symmetry axes highlighted in gray as pentagons, triangles and circles, respectively. **(c)** Pentameric capsomer viewed down the fivefold axis, and **(d)** hexameric capsomer viewed down the icosahedral-threefold/quasi-sixfold axis, showing OG labeling at K105, nearly within subunit interfaces.

Although the BMV coat protein contains 12 lysine residues, experiments have shown that approximately two dyes per chain tend to be covalently bound at saturation. ^5,6^ These lysines can be categorized into three distinct groups based on their structural location: interior, exterior, and interfacial. ^6^ This classification was achieved through experimental and computational analyses, which revealed that the reactivity of lysines is influenced by their accessibility, related to their position within the capsid architecture. Since lysines on the capsid interior (on the N-termini) are not relevant to SR, dyes bound at the more rigid, structurally integral sites were implicated as the active subset responsible for cooperative emission. Only K64, K105, K111, K165 of the exterior subset appear to be accessible for OG labeling. ^6^

Compared to other candidate labeling sites, K105 showed reduced accessibility at pH 4.5, where the VLP is at its most compact state. Importantly, plant viruses like BMV undergo a swelling transition in response to elevated pH or removal of divalent cations,^18^ and the dye conjugation reaction is carried out at pH 6.0, where the swollen state may increase K105 exposure.^6^ In native VLPs, K105 is located within a shallow depression, partially occluded by the capsid surface (Figure 1c-d), which restricts mobility of attached OG compared to other exterior lysines. The nearly interfacial character of position K105, which places spatial constraints on the motion of OG, may promote superradiance by limiting dephasing.

Here, all-atom molecular dynamics (MD) simulations on the microsecond timescale are used to examine a model SR particle, BMV-OG_K105_, as well as its structural scaffold, the native BMV capsid. Importantly, MD simulations of VLPs are uniquely suited to study the emergent properties of intact assemblies,^19^ such as the collective motions underlying fluorophore synchronization and SR in the BMV-OG_K105_ system. The present work focuses on the particle’s ground state dynamics, establishing a baseline for VLP biophysical characteristics at equilibrium, prior to excitation. Analyses reveal how scaffold behavior is influenced by chemical modification with OG, and how the geometry and fluctuations of the scaffold in turn lead to dye coupling relevant to cooperative emission. These molecular-level insights into the role of capsid architecture in promoting SR from a multi-fluorophore emitter are critical for establishing design rules for engineering SR-VLPs as novel fluorescent probes.

## Results

### Dye conjugation at K105 increases VLP volume

Both BMV and BMV-OG_K105_ exhibited an immediate sharp decrease in polyhedral shell volume upon the start of unrestrained dynamics, followed by a gradual recovery to the crystallographic ^17^ volume by 100 ns (Figure 2a). This response likely relates to low internal solvent density at simulation onset (Figure S1), coupled with low rates of water exchange across the capsid surface (Figure S2a-b), resulting in a protracted correction of the water disparity. The systems continued to expand afterward, but diverged at 200 ns, with BMV-OG_K105_ consistently maintaining a larger volume than BMV. Because the dyes are attached at K105, a residue located near inter-subunit interfaces with the potential to nestle between neighboring coat protein chains (Figure 1c-d), it appears that their presence may promote modest capsid expansion to accommodate the bulky, dianionic fluorescein moities within pentamers and hexamers. The volume gradient converged to *<*0.01% by the one microsecond mark, with VLPs ultimately reaching sizes of about 6.08% and 8.52% that of the crystal model used for initial coordinates (PDB 1JS9),^17^ for BMV and BMV-OG_K105_, respectively. The last 800 ns of simulation, following equilibration of internal cavities, structural relaxation (Z score *<* 2, Figure S3), and divergence of VLPs, was taken for subsequent analysis.

**Figure 2:**
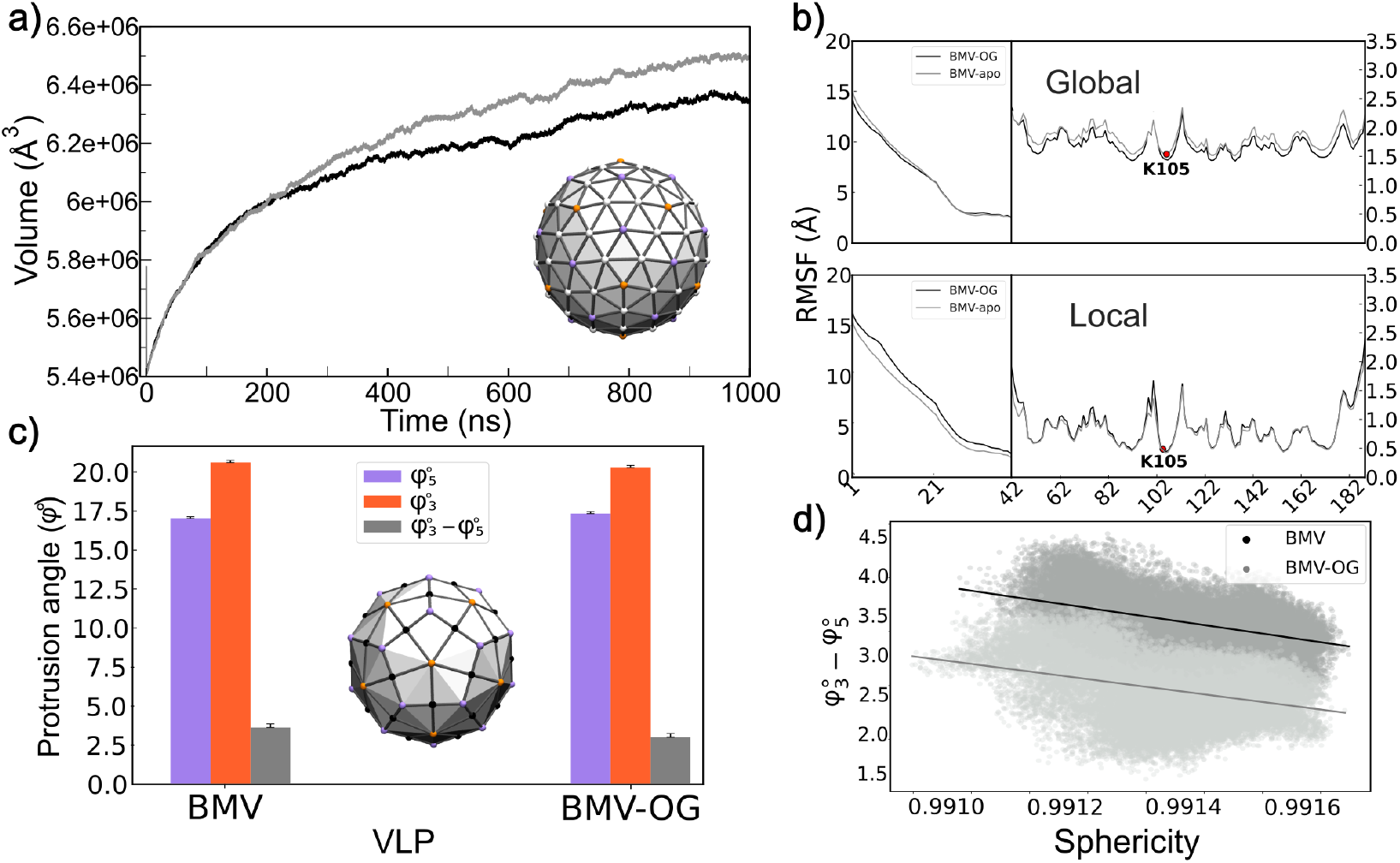
**(a)** Time series of shell volume for BMV and BMV-OG. Inset: polyhedron used for calculations. **(b)** Per-residue C*α* RMSF profiles comparing BMV and BMG-OG. Global alignment to the capsid (top) local alignment to individual chains (bottom). **(c)** Comparison of the average fivefold (*φ*_5_; purple) and threefold (*φ*_3_; orange) protrusion angles between BMV and BMV-OG, and their difference (*φ*_3_ − *φ*_5_ ; gray). Inset: kite-faced polyhedron with fivefold vertices in orange and threefold vertices in purple. **(d)** Correlation between faceting (*φ*_3_ *− φ*_5_) and sphericity, showing that greater faceting is associated with lower sphericity.

### Dye conjugation at K105 subtly influences VLP fluctuations

C*α* root-mean-square fluctuations (RMSF) of the coat protein indicate that the internal motion of VLP-incorporated chains is unaltered by labeling with OG at K105 (Figure 2b, local alignment). However, when considered in the context of the intact VLP, chains of the BMV-OG_K105_ particle exhibit subtly higher RMSF relative to BMV (Figure 2b, global alignment). This minimal increase captures spatial translations, suggesting greater inter-subunit flexibility when OG is attached. Such a result likely arises from expansion of BMV-OG_K105_ to accommodate dyes at K105, as reflected in its larger polyhedral shell volume. Notably, K105 remains among the least mobile residues, a property intrinsic to the native BMV capsid. Given that average K105 fluctuation in the VLP is only *∼*1.5 Å and scaffold rigidity is thought to be important for SR,^5^ this observation is likely functionally relevant.

### Dye conjugation at K105 minimally alters VLP morphology

A kite-faced polyhedron, including one kite per asymmetric unit, was used to characterize the shape of VLPs. Previous applications of this method revealed increased faceting of hepatitis B capsids in response to bound interfacial molecules, causing fivefold vertices to protrude and threefold vertices to flatten, resulting in a structure more closely resembling a geometric icosahedron.^20,21^ Remarkably, VLPs examined here exhibit the opposite behavior, both morphologically and in terms of ligand influence. OG attached at K105 leads to a less faceted capsid (Figure 2c). Further, faceting in the BMV system emphasizes pentagonal surfaces of the polyhedron rather than triangular. That is, fivefold vertices flatten and threefold vertices protrude, resulting in a structure more closely resembling a geometric dodecahedron, the icosahedral dual. Although the morphological disparity is minor, with the average overall faceting angle (*φ*_3_ *− φ*_5_) differenting by only *∼*1^*°*^, the adjustment likely again reflects alterations at subunit interfaces to sterically and electrostatically accommodate OG. Correlation analysis shows that increased faceting coincides with reduced sphericity (Figure 2d). While the sphericity of both VLPs decreased *<*0.1% relative to the crystal model used for initial coordinates (PDB 1JS9,^17^ Figure S4), both remain 99% sphere-like following relaxation while a true dodecahedron is only 91% sphere-like.

### Dye conjugation at K105 indirectly alters pore apertures

Given that capsid pores are located on the fivefold and threefold (quasi-sixfold) symmetry axes, morphological shifts involving these vertices could alter pore topology. Characterization of pore radii using HOLE^22^ indicate that pentameric pores are consistently narrower than hexameric pores, reflecting tighter geometric packing of five versus six protein chains within these subunits. In the case of BMV, pores exhibit median minimum diameters of 5.2 and 14.0 Å in pentamers and hexamers (Figure 3a-b), which readily accommodate 2.8-Å water molecules. Thus, the larger and more numerous hexameric pores are the primary channel for solvent exchange in the native VLP, with water crossing events between the particle interior and exterior occurring at influx and efflux rates of 1, 625 *±* 41 and 1, 613 *±* 41 molecules/ns (Figure S2a-b). In BMV-OG_K105_, pentameric pores expand by 9% while hexameric pores contract by 9% relative to BMV, representing median minimum diameters of 6.2 and 13.6 Å (Figure 3c-d). This shift results in an 8-9% increase in the frequency of water crossing events, with influx and efflux rates of 1, 763 *±* 46 and 1, 747 *±* 47 molecules/ns (Figure S2a-b).

**Figure 3:**
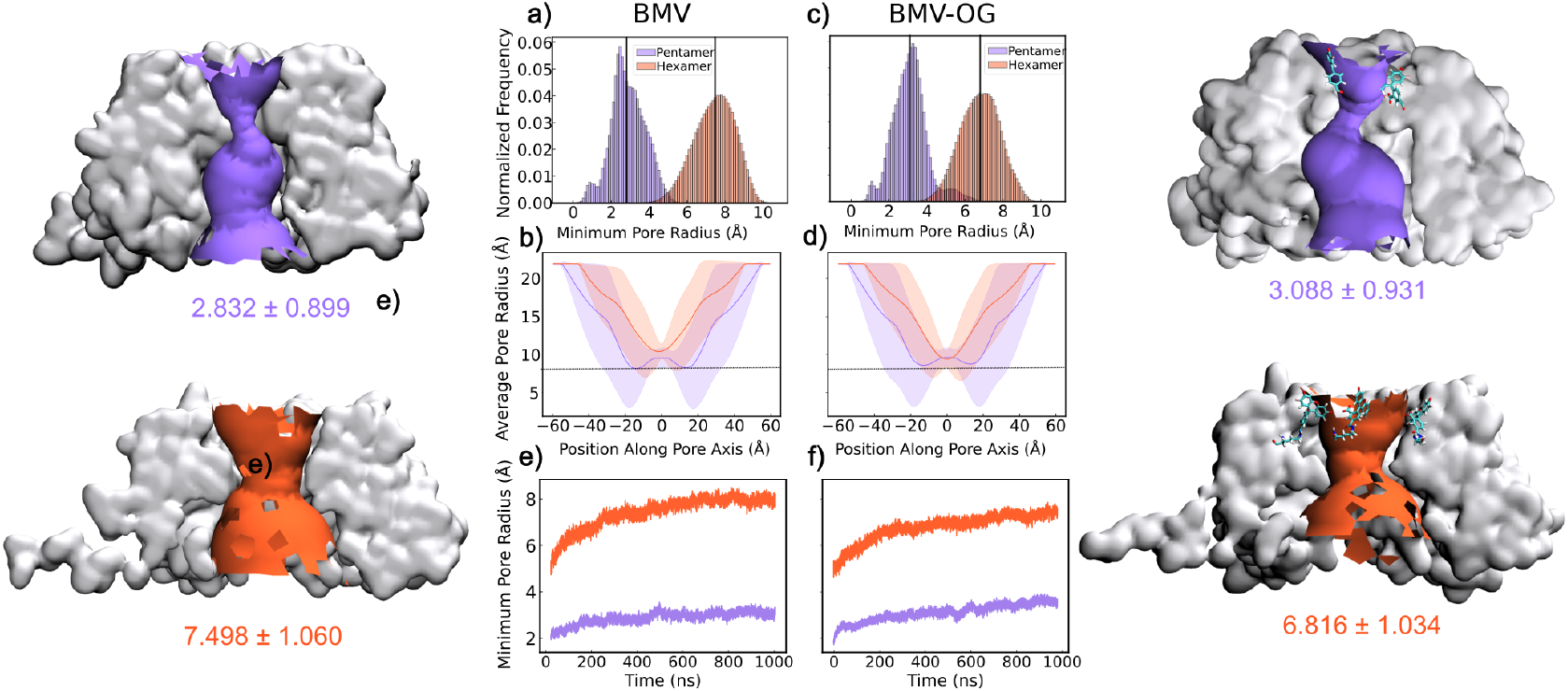
Characterization of pores in pentameric (purple) and hexameric (orange) capsomers for **(left, a-b)** BMV and **(right, c-d)** BMV-OG_K105_. Average radial profiles along the pore axes for **(a)** BMV and **(c)** BMV-OG_K105_. Solid lines indicate the median radius across all pores of a given capsomer type. Distributions of minimum pore radii for **(b)** BMV and **(d)** BMV-OG_K105_. Time evolution of average minimum pore radii for **(e)** BMV and **(f)** BMV-OG_K105_ Representative surface renderings of pores are shown for both VLPs, with OG dyes depicted as sticks in BMV-OG_K105_. Pore lining residues along the bottleneck of the pore aperture are S129, K130, E131. Median and standard deviation of corresponding distributions are indicated alongside structures.

The time evolution of pore diameters for both VLPs reveals meaningful changes compared to the experimental BMV structure, ^17^ including the ability to simultaneously accommodate 1-2 additional water molecules within the most narrow region of hexameric channels (Figure 3e-f), indicating that crystal structures can represent a poor basis for inferring capsid permeability. While pore topology is largely conserved despite OG conjugation at K105, the presence of the dianionic dye influences solvent exchange rates by interacting with the three residues that define the bottleneck of the pore opening: S129 and K130, as well as with E131 via an Mg^2+^-mediated contact. Thus, changes in pore dimensions do not arise from the dyes themselves defining the bottleneck, but from the dyes’ influence on nearby polar residues that dictate the size of the aperture. Interestingly, expansion of pentameric pores and contraction of hexameric pores when OG is bound correlates with the VLP exhibiting reduced flattening at fivefolds and reduced protrusion at threefolds compared to native BMV.

### BMV-based VLPs are selectively permeable to anions

Despite bidirectional water exchange across the capsid surface, both VLPs exhibit reduced and selective permeability with respect to ions. Cl^*−*^ influx and efflux rates of 11.0 *±* 0.2 and 11.3 *±* 0.2 molecules/ns for BMV compared to 9.7 *±* 0.2 and 9.9 *±* 0.2 molecules/ns for BMV-OG_K105_ reflect fewer crossing events in the presence of the dianionic dye (Figure S2c-d). Weak Cl^*−*^ occupancy is apparent at the mouth of capsomer pores near R103/K130 in BMV (Δ*G* = *−*1.0 kcal mol^*−*1^), but absent in BMV-OG_K105_ owing to the increased local charge of 10-12*e* (Figure 4a). Similar Cl^*−*^ occupancy along inter-capsomeric crevices, which are distal to K105-linked OG and lined by basic residues like arginine and lysine, remains unaffected. In contrast, Na^+^ permeates the capsid only rarely, with influx and efflux rates of 0.1 molecules/ns for both VLPs; no Mg^2+^ exchange was observed during simulations (Figure S2e-f). The crystal structure of BMV shows Mg^2+^ complexed at the quasi-threefold vertex as well as the fivefold pores^17^ (Figure S5), and although these were included in the MD initial model, classical ion parameters do no reproduce the coordination behavior of divalent cations. Nevertheless, the electrostatic nature of Mg^2+^ is sufficient to capture its binding with D127/E131 within the fivefold pores of both VLPs (Δ*G* = *−*8.5 kcal mol^*−*1^), consistent with crystallographic data (Figure S5).

**Figure 4:**
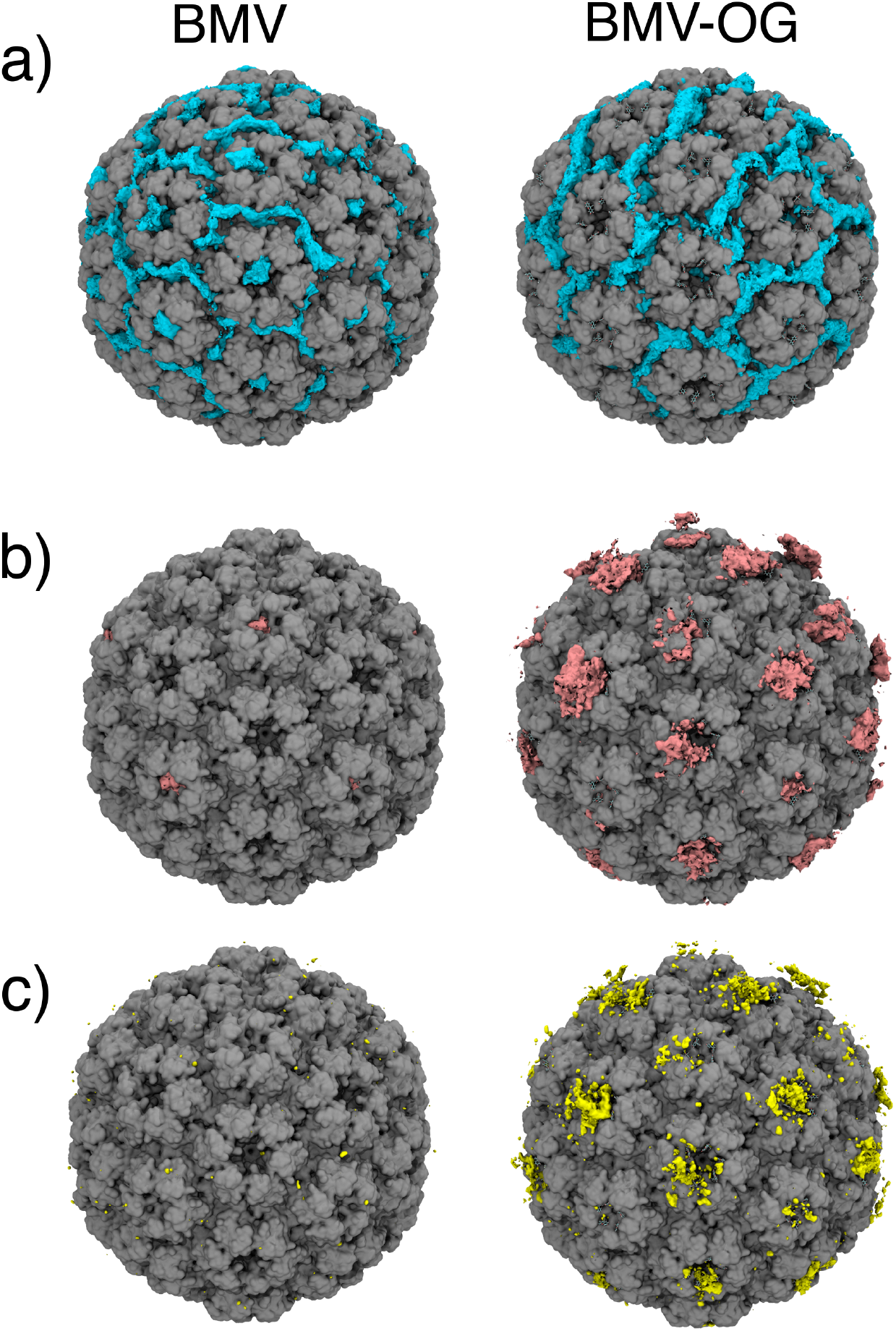
**(a)** Chloride (cyan), **(b)** magnesium (pink), and **(c)** sodium (yellow) ion localization maps are shown overlaid on the capsid surfaces for BMV (left) and BMV-OG (right). Occupancy maps were computed from simulation trajectories and rendered at a free energy threshold of Δ*G* = *−*1.0 kcal mol^*−*1^ to highlight favorable ion-binding regions.

Owing to the electronegativity of the dye, BMV-OG_K105_ displayed weak Mg^2+^ and Na^+^ localization externally, as a counterbalance over the dye-decorated capsomers (Δ*G* = *−*1.0 kcal mol^*−*1^), Figure 4b-c). Strong Mg^2+^ interactions with the pore-defining residues of VLP pentamers, but not hexamers, a characteristic of the native capsid (Figure S5), may impede water flux and contribute to the observed solvent exchange rates. Given the established role of divalent cations in capsid stabilization ^18^ and the importance of scaffold stability in promoting SR, ^5^ bound Mg^2+^ is likely an important contributor to an effective BMV-based SR-VLP.

### Pentamer-bound OG_K105_ exhibits dynamics most conducive to SR

Ground-state dynamics and orientation of VLP-conjugated dyes suggest how their mobility and relative alignment within capsomers could influence transition dipole coupling underlying SR. The transition dipole of OG is defined as shown in Figure 5a. Dyes located in pentamers (A chains) in BMV-OG_K105_ are more likely to display low RSMF values than those in hexamers (B and C chains), indicating that pentamer-bound OGs tend to be less mobile (Figure 5b). Dyes located in pentamers are also more likely to be situated in close proximity, rarely sampling the extreme inter-OG spacing observed in hexamers (Figure 5c). These 5-10 Å separation distances are within the range over which excitonic interactions can occur, ^23–25^ and demonstrate that pentamer-bound OGs are more frequently positioned to experience strong dipole-dipole coupling. Finally, as evidenced by the orientation factor *κ*^2^, dyes located in pentamers have a higher probability to adopt relative alignments that predispose them to cooperative emission (Figure 5d).

**Figure 5:**
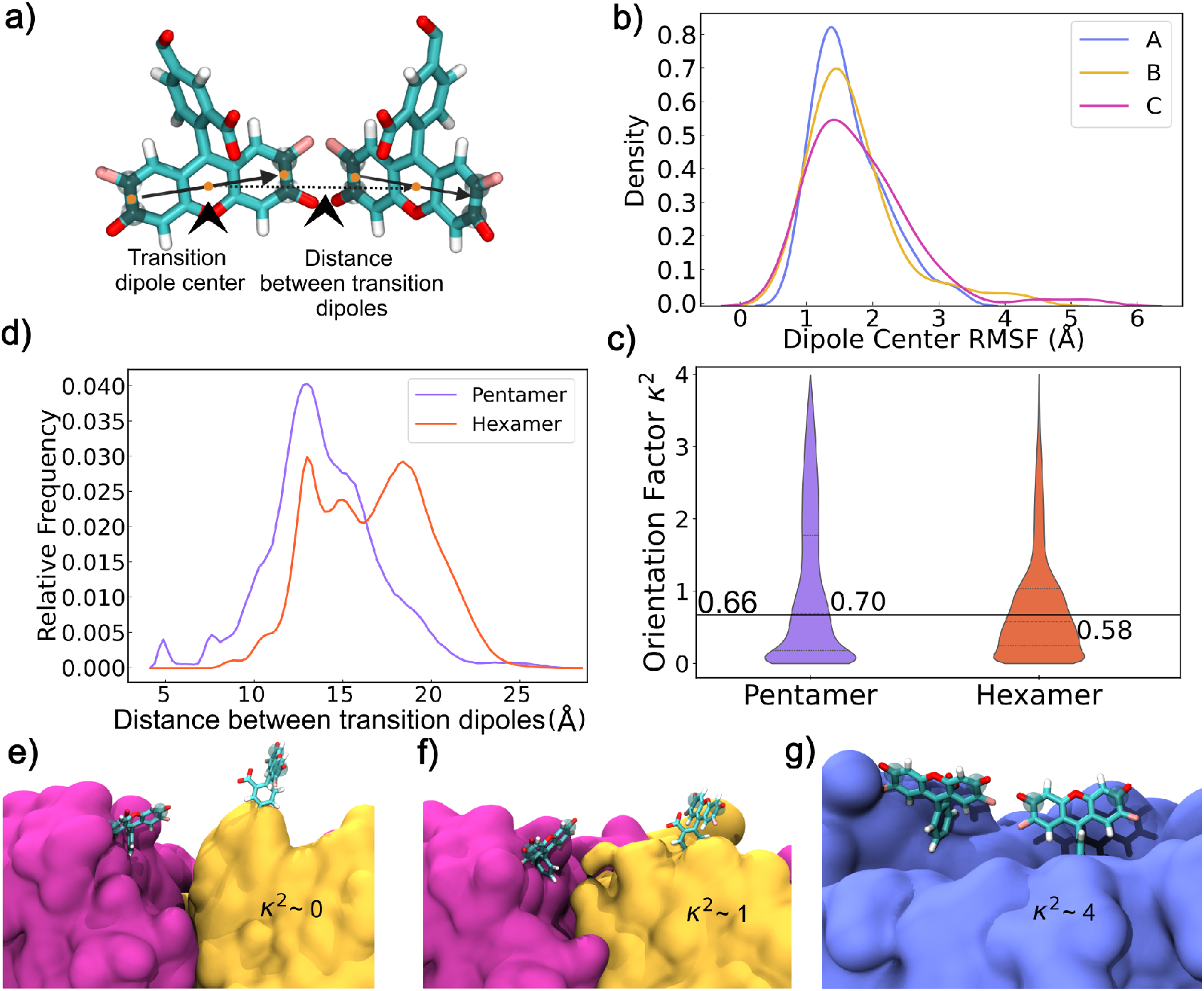
**(a)** Schematic of OG dyes showing atoms used to define the transition dipole vectors (highlighted atoms) and their midpoints used to compute dipole centers and inter-dye distances. **(b)** Distribution of root-mean-square fluctuations (RMSF) of dipole centers for dyes attached to quasi-equivalent A, B, and C chains. **(c)** Violin plots of the orientation factor (*κ*^2^) for pentamer-and hexamer-bound dye pairs. Horizontal line indicates the isotropic average value of 2*/*3 or 0.66. **(d)** Distribution of inter-dye distances measured between transition dipole centers in pentamers and hexamers. **(e)** Representative snapshots of dye pairs with different *κ*^2^ values, illustrating **(e)** orthogonal (*κ*^2^ *∼* 0), **(f)** perpendicular (*κ*^2^ *∼* 1), and **(g)** nearly parallel (*κ*^2^ *∼* 4) orientations.

While *κ*^2^ is classically used in Förster resonance energy transfer (FRET) to describe transfer efficiency, in the context of SR it captures the propensity for excitonic coupling between emitters. ^26,27^ *κ*^2^ spans the continuous set [0-4],^28^ and pentamer- and hexamer-bound OGs explore the full range of values in the ground state. Dyes attached to both capsomers most frequently sample *κ*^2^ *∼* 0, corresponding to orthogonal or out-of-phase orientations that prevent cooperative emission (Figure 5d,e). Alignments associated with *κ*^2^ *∼* 1 are also common, corresponding to a mutual orientation of dipoles at the magic angle (54.7^*?*^) where energy transfer is non-radiative (Figure 5d,f). Notably, the distribution median for both pentamer- and hexamer-bound OGs is close to *κ*^2^ *∼* 2*/*3, corresponding to transition dipoles rapidly and uniformly sampling all possible relative orientations ^28^ (Figure 5d). However, dyes located in pentamers exhibit a broader distribution tail extending toward *κ*^2^ *∼* 4, corresponding to nearly parallel and in-phase transition dipoles conducive to SR (Figure 5d,g). Thus, in BMV-OG_K105_, the pentameric capsomer environment promotes more orientational anisotropy, while the hexameric environment allows greater orientational disorder. That pentamer-bound OGs have a higher likelihood of producing SR-ready dye alignments even prior to excitation highlights both the role of capsid geometry and the nearly interfacial position of K105 in modulating SR emission from BMV-based VLPs.

## Discussion

Microsecond all-atom MD simulations of BMV-based VLPs in the ground state reveal that conjugated OG induces only subtle structural adjustments in the capsid scaffold. These findings are consistent with experimental evidence that dye-decorated BMV is morphologically indistinguishable from native virions.^6,14^ However, even subtle alterations in structure can affect particle dynamics. Small changes at inter-subunit interfaces in response to accommodating bulky, negatively charged OG lead to a discernable increase in global VLP fluctuations. Since lysine modification with S-methylthioacetimidate (SMTA) was shown to decrease apparent melting temperature at low pH, suggesting mild loosening of the BMV interfaces,^14^ larger OG might be expected to have a more pronounced impact. On the other hand, the fluorescein moiety of OG provides favorable electrostatic contacts that bridge neighboring protein chains when attached at K105, which may mitigate adverse steric effects. Since authentic SR-VLPs include OG on other external lysines (*e*.*g*., K64, K111, K165), interactions of these additional dyes with the capsid surface may further account for the structural and chemical stability of the particles according to experiments. ^5,6^

Superradiant emission has been observed to diminish in the presence of any factors that increase structural disorder in the scaffold.^5^ For example, buffer conditions including low pH and the presence of divalent cations play a central role in maintaining the compact, ordered VLP architecture necessary for coherent emission. BMV and closely related viruses like cowpea chlorotic mottle virus (CCMV) undergo a swelling transition, arising from electrostatic repulsion at the quasi-threefold vertices in response to increased pH and/or removal of divalent cations. ^18,29^ All-atom MD simulations employed here used protonation states consistent with pH 4.5, as well as explicit solvent/ions to represent SAMA buffer and maintain the compact, unswollen state of the capsid scaffold. While the BMV crystal structure captures stabilizing Mg^2+^ bound at the fivefolds and quasi-threefolds (the junctures of pentamers with two adjacent hexamers),^17^ free energies of ion localization from MD indicate Mg^2+^ binding only within the fivefolds. Although classical fixed-charge models do not reproduce the coordination behavior of divalent cations, Mg^2+^ does not exhibit typical octahedral coordination geometry in the BMV crystal^17^ and appears to act primarily as a counterion bridging a cluster of charged residues. Despite the absence of strong Mg^2+^ occupancy in the quasi-threefolds in MD, the volume expansion of VLPs does not reflect the structural changes associated with capsid swelling, which would be apparent on the microsecond timescale. ^30,31^

These observations confirm that the buffer conditions mimicked by the simulation are sufficient for stabilizing the unswollen state of the capsid appropriate for low pH. Importantly, classical MD represents pH in terms of fixed protonation states of titratable residues, which may be assigned based on the local environment of the starting structure, ^32,33^ but are not updated during dynamics. MD captures strong Mg^2+^ occupancy within pentameric pores at fivefolds, where the five copies of E131 were predicted to be deprotonated and negatively charged at pH 4.5 (Figure S5a-b). Similarly, Mg^2+^ occupancy is not observed within hexameric pores at threefolds, where the six copies of E131 were predicted to be protonated and neutral, owing to being spaced much farther apart than in pentamers (Figure S5c-d). With respect to the three copies each of E84/D148 at pseudo-threefolds, 5 out of 6 (83%) residues were predicted to be protonated and neutral, explaining the absence of Mg^2+^ occupancy during MD (Figure S5e-f). Given that local sampling suggests 2 out of 6 (33%) residues would be consistently protonated (Figure S6), along with the reduced likelihood of the acid forms at increased pH, the presence of pseudo-threefold Mg^2+^ in the pH 5.2 crystal may report on the true charge state of E84/D148. Clearly, bound Mg^2+^ counterbalances this cluster of residues when deprotonated, allowing the particle to maintain its compact, unswollen state. Altogether, these results support a purely electrostatic role for divalent cations in stabilizing BMV, rather than serving as chemical coordination centers.

Perhaps the most interesting insight from MD simulations is how the capsid scaffold affects the behavior of the dyes, particularly their orientation with respect to each other. Both the spatial distribution and dynamics of dyes conjugated on the VLP dictate the extent of excitonic coupling that underlies collective emission in SR. In BMV-OG_K105_, the quasi-equivalence of BMV subunits creates distinct local environments for OG. Pentamer-bound dyes experience somewhat less conformational freedom than hexamer-bound dyes, owing to more persistent protein-dye interactions. The closer packing of A chains enables dyes to situate closer together and increases the probability of parallel dipole alignment, a pre-requisite for coherent emission. In contrast, hexamer-bound dyes display random orientations more frequently, suggesting weaker excitonic coupling on average. From an SR-VLP design perspective, the combination of restricted mobility, shorter inter-dye spacing, and ordered dipole alignment makes pentamer-bound dyes especially well-suited to act as cooperative emitters capable of SR, especially when OG is attached to K105, the most rigid lysine in BMV. However, radiation brightening under pulsed excitation only occurs with dye loadings exceeding 170-200,^5^ such that the 60 pentameric K105 positions would alone be insufficient for SR. While capsid-wide saturation of K105 results in a total of 180 conjugated OGs consistent with the SR-active loading threshold, experiments demonstrate that other external lysines (*e*.*g*., K64, K111, K165) are also commonly labeled in authentic BMV-based VLPs.

## Conclusion

This work highlights the power of all-atom MD simulations in connecting structural biophysics with emergent optical phenomena. While experimental techniques can individually reveal static structures or bulk optical outputs, MD provides a dynamic, atomistic perspective linking the two. The present results show how dye conjugation can reshape intact capsid properties in ways that propagate down to local environments, ultimately influencing dye orientation and coupling that promotes SR. Characterization of BMV-based VLPs in their ground state establishes an essential baseline for future work investigating particle dynamics in the excited state. Finally, beyond the mechanisms underlying SR imaging probes, these findings are relevant to understanding the nature of the undecorated scaffold, the BMV capsid, which plays essential functional roles during the pathogen’s infection in plants.

## Methods

### Model Construction

An all-atom model of the BMV capsid was generated using crystal structure PDB 1JS9, ^17^ which was selected for its resolution (3.4 Å) and relative completeness compared to other available experimental structures. Notably, when aligned on residues 49-176 to exclude the disordered/incomplete termini, 1JS9 and alternative cryoEM-derived structures 3J7L^34^ and 6VOC^35^ have a total RMSD less than 1.0 Å. Missing residues 1-40 of the A chain and 1-24 of the B chain in 1JS9 were reconstructed as previously described^36^ to produce a model of the full-length, intact T=3 capsid.

An analogous model of the superradiant virus-like particle (BMV-OG) was generated by labeling the BMV capsid with Oregon Green 488 (OG). While experiments indicate that OG can be conjugated to K64, K105, K165, and other lysines on the N-terminus, ^36^ K105 appears to be most relevant for promoting superradiant emission. Thus, the model investigated here was saturated with K105-linked OG to produce a symmetric and homogeneously-labeled BMV-OG incorporating 180 copies of the fluorophore. Manual adjustment of K105 to the g+/t/g+ rotamer, which is freely sampled in the unlabeled BMV capsid, was required to sterically accommodate OG on all three quasi-equivalent chains.

PDB2PQR^32,33^ was used to assign protonation states to titratable residues appropriate for pH 4.5, representing the unswollen state of the capsid. Local Na^+^ and Cl^*−*^ ions were placed using the cionize plugin in VMD,^37^ maintaining all crystallographic Mg^2+^ ions.^17^ The BMV and BMV-OG models were immersed in 322.5-Å^3^ solvent boxes containing 50 mM NaCl and 8 mM MgCl_2_ to mimic SAMA buffer, consistent with environmental conditions used in experimental studies of BMV-OG. ^5^ The final systems encompassed 3.25 million atoms.

### Dye Parameterization

#### Bonded Parameters

The OG core was parameterized based on analogy to the Alexa Fluor family from the AMBER-DYES^38^ force field. Substituents of the conjugated ring system were parameterized based on analogy from the GAFF2 generalized AMBER force field.^39^ Appropriate matches were identified for every atom type. Parameters for the lysine linkage were taken from AMBER-DYES.

#### Nonbonded parameters

The OG structure was capped with N-methyl amide and the geometry was optimized in vacuum at the HF/6-31G* level of theory with Gaussian16. ^40^ Charges were computed at the B3LYP/6-31G* level of theory based on grids of ten concentric layers with a total number of 10.5 *×* 10^4^ grid points. A restrained electrostatic potential (RESP)^41^ charge fit was carried out using a weight of 0.0005 for all heavy atoms. The second stage fit typically employed with the RESP protocol was unnecessary owing to the absence of aliphatic/non-aromatic carbons in OG. Consistent with AMBER-DYES, Lennard-Jones parameters were taken from GAFF2.

#### Ensemble-fitted charges

The optimized OG structure was immersed in a box of TIP3P^42^ water with a 20-Å solvent buffer in each dimension. Using the preliminary RESP charges, a 10-ns MD simulation was performed with AMBER18^43^ in the isothermal-isobaric (NPT) ensemble. The Langevin thermostat and Monte Carlo barostat were invoked to maintain target conditions of 300 K and 1 bar. A simulation timestep of 1 fs was employed without bond constraints, along with a nonbonded cutoff of 14 Å. Frames were extracted from the trajectory at 1-ns intervals to serve as starting configurations for ten independent simulations performed in the microcanonical (NVE) ensemble. Each system was equilibrated for 50 fs, followed by 200 fs of production sampling. The AMBER18 quantum mechanics/molecular mechanics (QM/MM) interface calling Gaussian16 was used to compute charges for OG for every step at the B3LYP/6-31G*//TIP3P level of theory. Final RESP charges were obtained by fitting to the aggregate production ensemble of 2,000 configurations.

### Molecular Dynamics Simulations

Simulations of the intact BMV and BMV-OG capsids were performed with NAMD3. ^44,45^ Simulation files were constructed using psfgen with the AMBERff-in-NAMD implementation,^46^ applying the ff14SB^47^ protein force field and TIP3P^42^ water model, along with custom OG parameters developed as described above. The models were subjected to energy minimization for 20,000 steps using the conjugate gradient algorithm. The models were heated from 50 K to 300 K over an interval of 5 ns, and backbone restraints of 5 kcal/mol were gradually removed over an additional interval of 5 ns. Conformational sampling was collected in the isothermal-isobaric (NPT) ensemble on the microsecond timescale.

Simulations used the r-RESPA integrator with a time step of 2.0 fs. Bonds to hydrogen were constrained with the SHAKE (solute) and SETTLE (water) algorithms. To most accurately apply AMBERff parameters, 1-4 interactions were scaled by 2.0 (non-bonded, SCNB) and 1.2 (electrostatics, SCEE), respectively, and Coulomb’s constant was set to 332.0522173 kcal/mol Å e^*−*2^ to match that of the native AMBER MD engine.^46^ Beyond a cutoff of 8 Å, long-range interactions were treated with a Lennard-Jones correction, and electrostatics were computed with Particle Mesh Ewald using a grid spacing of 1.5 Å and eighth-order interpolation. Full electrostatic evaluations were performed every other time step. System temperature of 300 K was maintained using the Langevin thermostat with a damping coefficient of 1 ps^*−*1^. System pressure of 1 bar was maintained using the Nosé-Hoover Langevin piston barostat, allowing isotropic cell scaling, with a piston oscillation period of 2,000 fs and damping timescale of 1,000 fs. Trajectory frames were saved every 10 ps.

### Trajectory Analysis

#### Capsid Morphology

To evaluate the shape of the BMV and BMV-OG capsids, polyhedrons were fitted to the BMV and BMV-OG capsids, as previously described.^48^ The polyhedron used to estimate particle sphericity included 180 triangular faces, one for each constituent protein chain (Figure 5). The surface area *SA* of the polyhedron was calculated as the sum over the areas of all triangular faces:

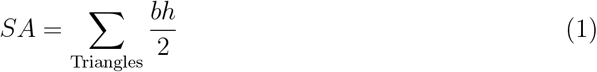

where *b* is the base length of each triangle and *h* is its height.

The volume *V* of the polyhedron was calculated as the sum over the volumes of triangular pyramids formed by each triangular face and the polyhedron centroid:

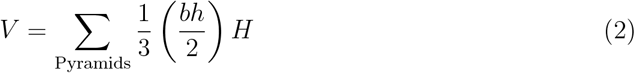

where *H* is the height of each pyramid.

The sphericity Ψ was calculated using the ratio of volume to surface area:

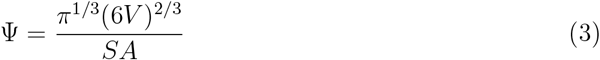

A value of Ψ = 1 corresponds to a sphere, while 0.911 corresponds to a dodecahedron.

The polyhedron used to estimate particle faceting included 60 kite-shaped faces, one for each asymmetric unit of the capsid (Figure 1c). Angles *φ*_5_ and *φ*_3_ were calculated as the orientation of the kite plane normal with respect to the fivefold and threefold symmetry axes, respectively. The faceting angle is defined as the difference *φ*_3_ *−φ*_5_, where larger values correspond to surface protrusion at threefold vertices and flattening at fivefold vertices, consistent with a less spherical, more icosahedral morphology.

#### Dye Orientation

The orientation factor *κ*^2^ was computed between neighboring pairs of OG dyes within each capsomer, for every 20 ps of the simulation. The transition dipole was defined as the vector connecting the midpoints between atom pairs (Figure 5a). This vector approximates the long-axis of the three-ring fluorescein moiety of OG and reflects the direction of its dominant electronic transition. Given 12 pentamers and 20 hexamers in the capsid, 60 (12 *×* 5) pentameric and 120 (20 *×* 6) hexameric neighboring OG pairs were evaluated, for a total of 180 unique dye-dye relationships analyzed per frame.

The orientation factor *κ*^2^ for neighboring OG pairs was computed using the standard Förster resonance energy transfer (FRET) equation:

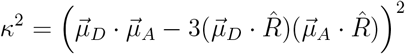

where 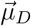 and 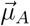 are unit vectors representing the donor and acceptor dipole moments, and 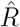 is the unit vector pointing from the donor center to the acceptor center.

The midpoint of the transition dipole vectors was used to represent the transition dipole centers. The proximity of neighboring OG pairs within a capsomer was computed as the distance between transition dipole centers. These distances provide a complementary geometric measure of dye packing, and were calculated every 20 ps of the trajectory.

### Root Mean Square Fluctuation (RMSF)

Root Mean Square Fluctuation (RMSF) is defined as the root mean-square-average distance between an atom and its average position over time, and provides a measure of its mobility. In this study, C*α*-RMSF was used to characterize both global capsid dynamics and local flexibility of the BMV coat protein, averaged across 180 copies. Structural alignments used the C*α* trace of the capsid crystal^17^ as a reference, excluding residues 1–41 to remove contributions from the flexible N-terminal tails. For assessing global dynamics, alignment was based on the entire capsid. To isolate local motions independent of inter-subunit displacements, alignment was based on individual protein chains. RMSF was calculated per residue and averaged across 180 copies. The mobility of OG with respect to the protein was characterized as RMSF of its transition dipole center, using the same local protein alignment protocol described above.

### Ion Localization

To characterize the spatial distribution of ions around BMV and BMV-OG capsids, 3D occupancy maps were calculated. Each trajectory was aligned to the capsid’s C*α* trace using the crystal structure ^17^ as a reference, and time-averaged maps for individual ion species (Na^+^, Cl^*−*^, Mg^2+^) were calculated using the <monospace>volmap</monospace> tool in VMD,^37^ with a grid spacing of 1.0 Å. Occupancies were based on 50,000 frames per trajectory using an interval of 20 ps. Occupancy values at each voxel, 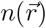, were converted to potentials of mean force (PMFs), representing local Gibbs free energy (Δ*G*) landscapes, using the following equation:

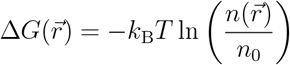

where *k*_B_ is the Boltzmann constant (0.008314 kJ/mol*·*K), *T* is the simulation temperature (300 K), and *n*_0_ is the mean ion occupancy in a reference bulk-like region at the capsid center. Separate values of *n*_0_ were determined for each ion species.

### Solvent Exchange

Water and ion exchange rates were computed using the fuzzy boundary implementation VMD’s `volinterior` to differentiate between the interior and exterior of the BMV capsid. ^49^ The QuickSurf representation of the protein molecular surface was defined with a radius scale of 1.8 Å, an isovalue of 1.6 Å, and a grid spacing of 1.0 Å. The intrinsically disordered N-terminal region (residues 1–41) was excluded from the surface because its extension into the capsid interior caused `volinterior` to misclassify nearby voxels as boundary, leading to underestimation of the internal volume. A total of 32 rays were cast from each voxel. Voxels with normalized occupancy greater than 0.95 were classified as interior; those between 0 and 0.85 were considered as exterior. Values between representing space within capsid pores and directly interacting with the VLP surface were excluded in order to track exchange between bulk solvent regions. Water and ion exchange events were identified by tracking atomic indices over time: if a molecule classified as exterior at reference frame *t*_0_ was found interior at *t > t*_0_, it was recorded as an inward crossing, and vice versa. Exchange events were evaluated every 5 ns within each 25 ns analysis block, with the reference time *t*_0_ reset at the start of each block to isolate short-timescale exchange dynamics. The average slope of the cumulative exchange profiles across all blocks was used to estimate the mean exchange rate, providing a robust measure of molecular flux across the capsid boundary (Figure S2).

## Abbreviations

BMV: Brome mosaic virus
MD: Molecular dynamics
OG: Oregon Green 488
QM/MM: Quantum mechanics /molecular mechanics
SR: Superradiance
VLP: Virus-like particle

## Author Contributions

J.A.H.-P. designed the project, parameterized OG, constructed models, and ran MD simulations. A.K. carried out large-scale MD trajectory analysis and developed data presentations. A.B. performed additional calculations to support interpretation of results. J.A.H.-P. and A.K. wrote the manuscript.

## Acknowledgement

This work was funded by NSF award CBET-2232718 to J.A.H.-P. This work was supported by the Delaware Advanced Research Workforce and Innovation Network (DARWIN), funded by NSF award OAC-1919839, and the BioStore resource, made possible by NIH through the Delaware IDeA Network of Biomedical Research Excellence, awards P20GM103446 and S10OD028725. Computer time on Delta at the National Center for Supercomputing Applications at the University of Illinois at Urbana-Champaign and on Anvil at the Rosen Center for Advanced Computing at Purdue University was provided by allocations BIO-230157 and BIO-240029 from the Advanced Cyberinfrastructure Coordination Ecosystem: Services & Support (ACCESS) program. ACCESS is funded by NSF awards #2138259, #2138286, #2138307, #2137603, and #2138296.

## Supporting Information

**Figure S1:**
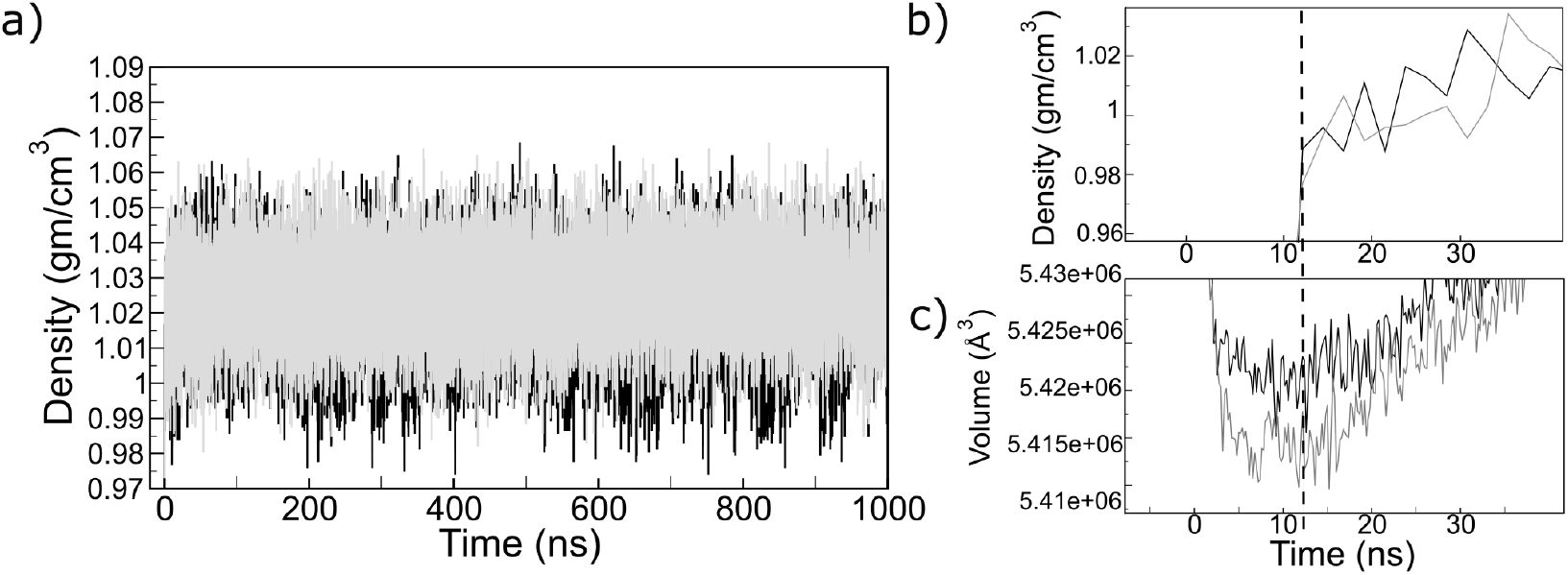
Correlation between internal water density and capsid volume. (a) Time evolution of water density within a spherical region of radius 20 Å centered inside the capsid for BMV (black) and BMV-OG (gray) over the 1 *µ*s trajectory. Both systems exhibit stable internal hydration densities consistent with bulk water (b, c) Early-time comparison of internal water density (b) and total capsid volume (c) reveals dips in both properties during the initial equilibration phase (vertical dashed line), indicating transient solvent depletion accompanying capsid compaction.

**Figure S2:**
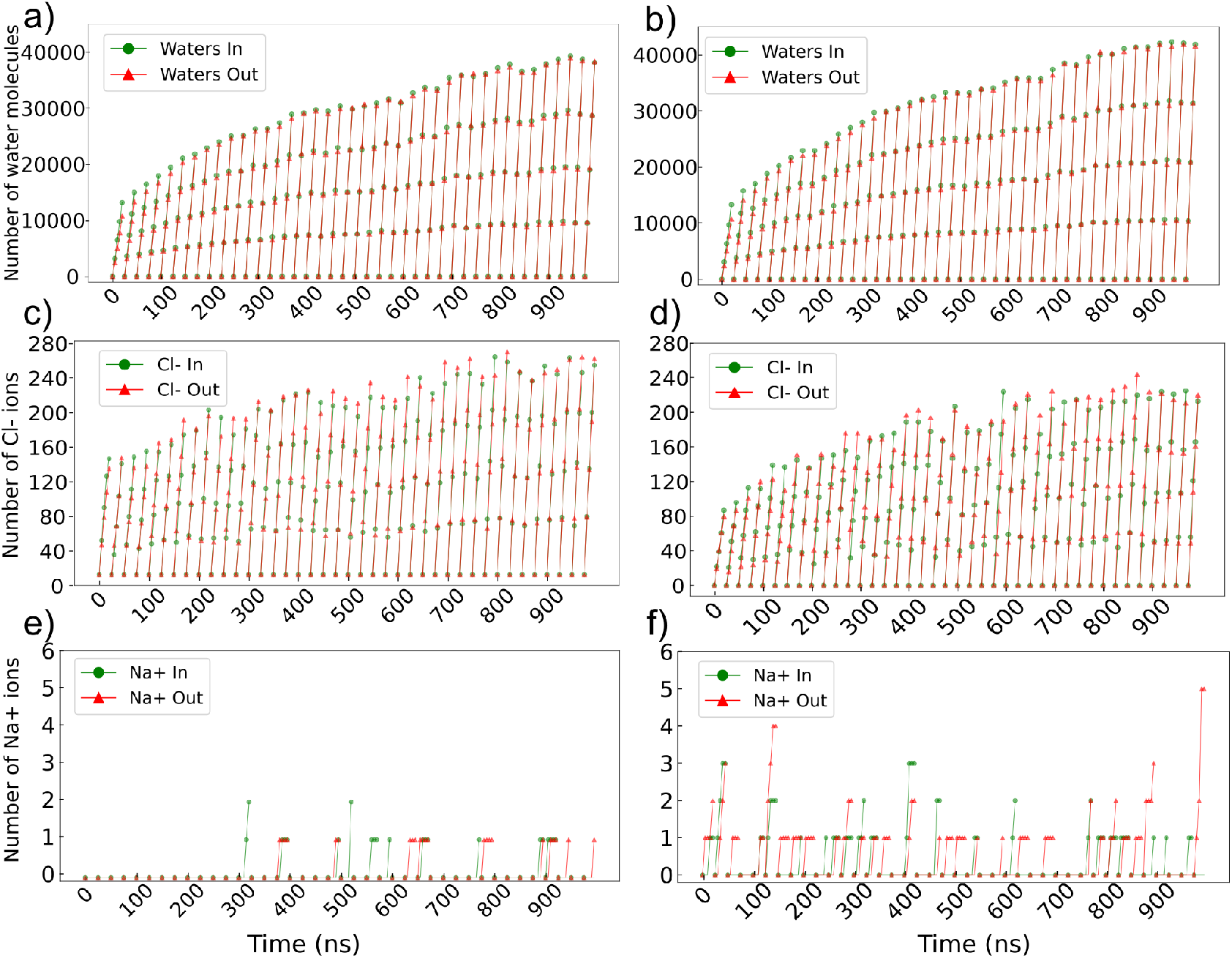
Cumulative numbers of solvent species moving inward (green) and outward (red) across the capsid surface are plotted versus simulation time, and the slopes of linear fits yield the exchange rates (reported as averages ± standard deviations). **(a–c)** BMV; **(d–f)** BMV-OG. **(a**,**d)** Water molecules exchange at rates of 1625 ± 41 ns^*−*1^ inward and 1613 ± 41.1 ns^*−*1^ outward in BMV, and 1763.1 ± 46.2 ns^*−*1^ inward and 1746.7 ± 47.1 ns^*−*1^ outward in BMV-OG. **(b**,**e)** Chloride ions exchange at 11.0 ± 0.2 ns^*−*1^ inward and 11.3 ± 0.2 ns^*−*1^ outward in BMV, and 9.7 ± 0.2 ns^*−*1^ inward and 9.9 ± 0.2 ns^*−*1^ outward in BMV-OG. **(c**,**f)** Sodium ions exchange rarely, with average rates of 0.1 ns^*−*1^ in both directions for both capsids.

**Figure S3:**
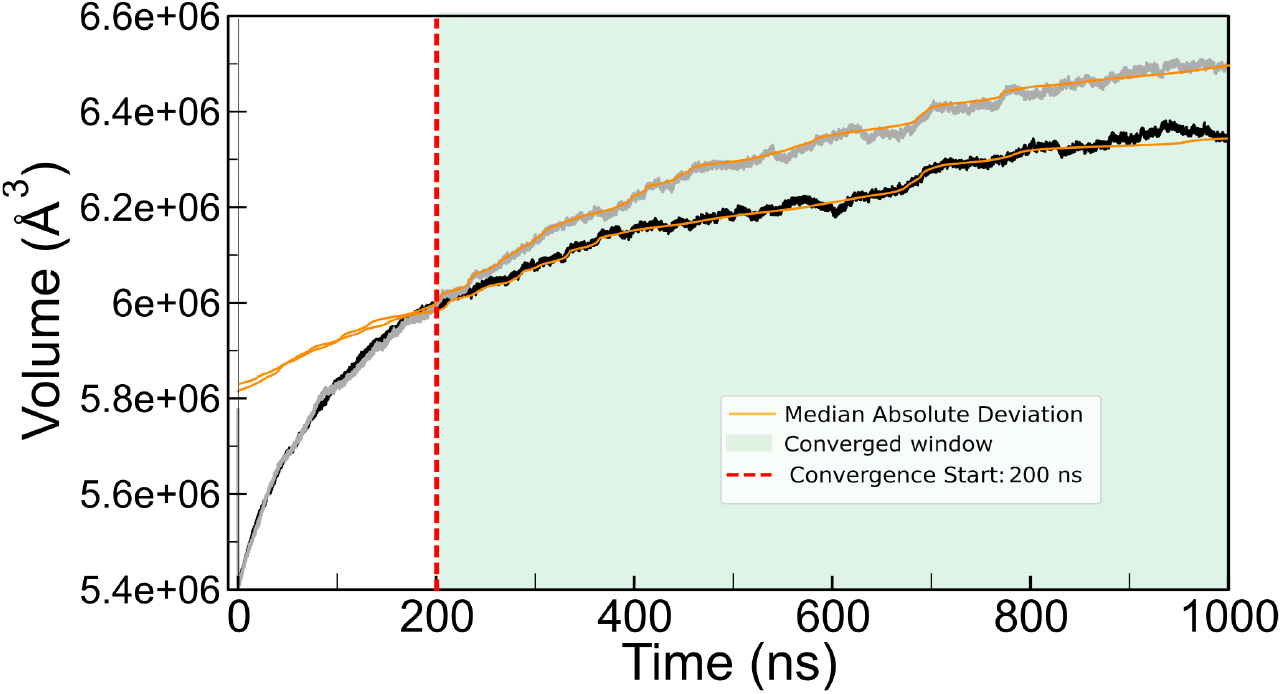
The rolling median and Median Absolute Deviation (MAD, orange) were used to identify the onset of convergence. The MAD quantifies deviations from the median and is related to the standard deviation of a normal distribution by *σ* ≈ 1.4826 × MAD, or equivalently, MAD ≈ 0.6745*σ*. The modified Z-score was calculated as the deviation of each data point from the rolling median, normalized by the MAD. Convergence was defined as the time point beyond which the modified Z-score remained within ±2 for at least 20000 consecutive frames, indicating that volume fluctuations had stabilized within two robust standard deviations of the median. A Z-score threshold of 2 was used to define the convergence window indicating that both capsids reached stable volume fluctuations after approximately 200 *∼* ns (red dashed line).

**Figure S4:**
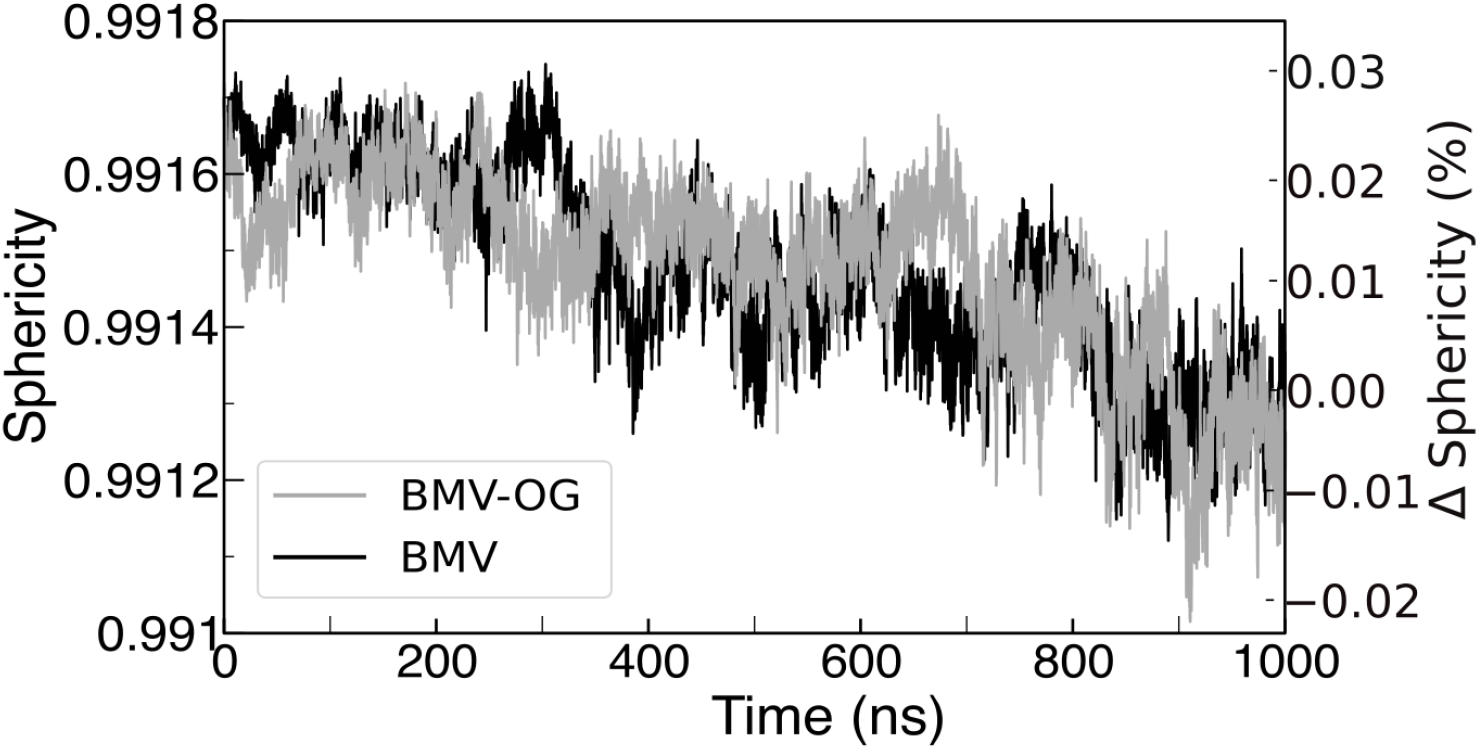
Comparison of sphericity for BMV (black) and BMV-OG_K105_ (gray) over 1 *µ*s of simulation time. The right axis shows the relative change in sphericity expressed as a percentage deviation from the initial value.

**Figure S5:**
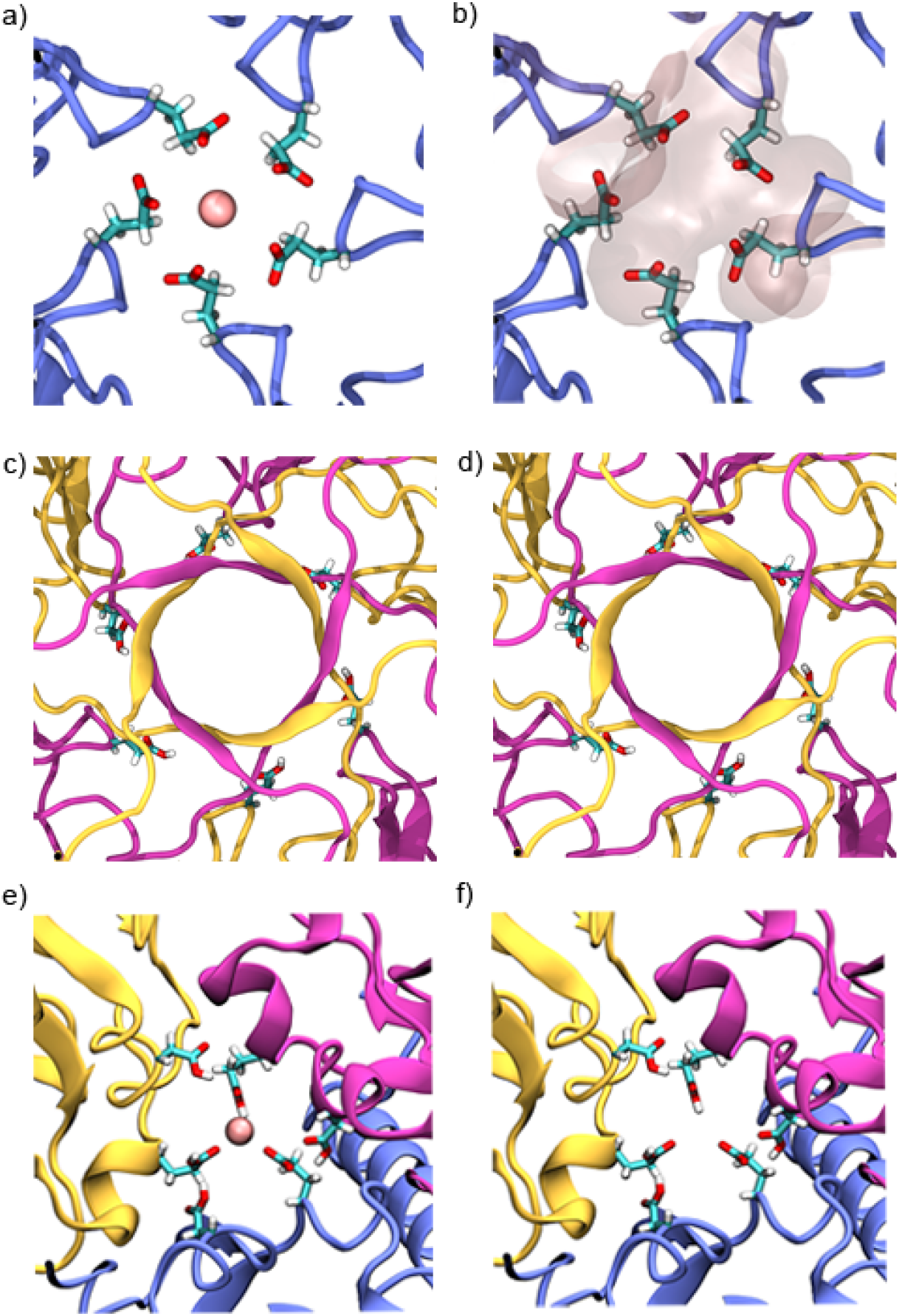
Mg^2+^ localization at pH 4.5. Fivefold vertex within pentamer showing E131 with **(a)** Mg^2+^ bound as in the crystal, versus **(b)** strong Mg^2+^ occupancy during simulation (Δ*G* = −9.7 kcal mol^*−*1^). PDB2PQR predicts E131 is deprotonated on all three chains. Threefold vertex within hexamer showing E131 with Mg^2+^ absent as in **(c)** the crystal and **(d)** lacking meaningful occupancy during simulation. PDB2PQR predicts E131 is protonated on all three chains. Pseudo-threefold vertex showing E84 and D148 with **(e)** Mg^2+^ bound as in the crystal, versus **(f)** the absence of Mg^2+^ occupancy during simulation. PDB2PQR predicts E84 is deprotonated (carboxylate form) on chains A and B but protonated on chain C, whereas D148 is protonated (carboxylic acid form) on all three chains.

**Figure S6:**
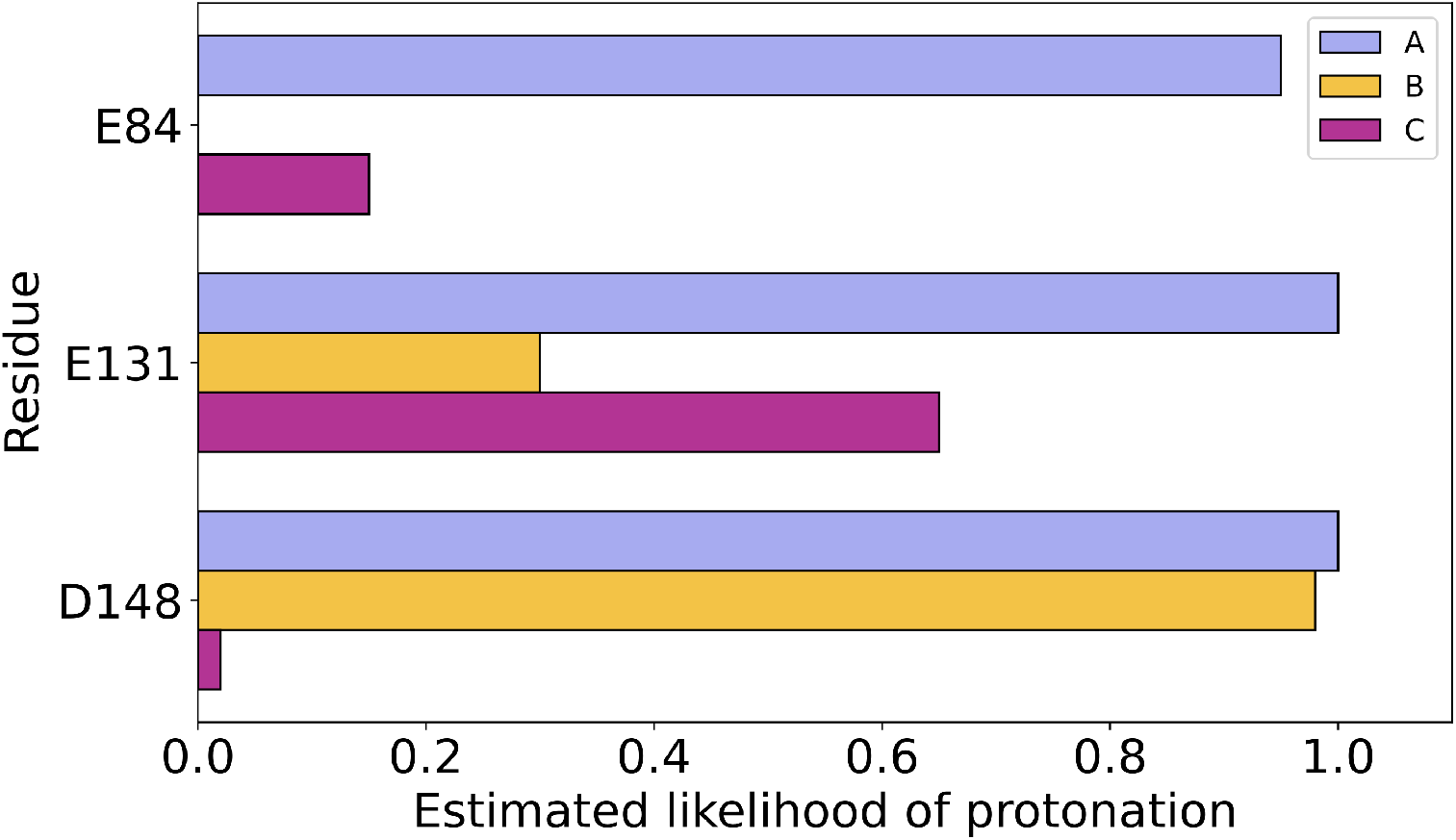
Estimated likelihood of protonation for residues E84, E131, and D148 across the three quasi-equivalent chains (A, B, C) in the asymmetric unit at pH 4.5. Calculation based on PDB2PQR predictions for a sequence of MD trajectory frames extracted from 30 ns simulation of the intact BMV capsid.^36^

